# Olfaction in Tephritidae: a balance between detection and discrimination

**DOI:** 10.1101/2024.03.14.584788

**Authors:** Gaëlle Ramiaranjatovo, Maud Charlery de la Masselière, Teun Dekker, Pierre-François Duyck, Sebastian Larsson Herrera, Bernard Reynaud, Vincent Jacob

**Affiliations:** CIRAD, UMR PVBMT, F-97410 St Pierre, La Réunion, France; Université de La Réunion, UMR PVBMT, F-97410 St Pierre, La Réunion, France; Chemical Ecology Unit, Department of Plant Protection Biology, Swedish University of Agricultural Sciences, Box 102, SE-230 53, Alnarp, Sweden; CIRAD, UMR PVBMT, F-98848 Nouméa, New Caledonia; IAC, Equipe ARBOREAL, BP 98857, Nouméa, New Caledonia; Hushållningssällskapet Skåne, Box 9084, 291 09 Kristianstad

**Keywords:** *Bactrocera dorsalis*, polyphagy, Fruit fly, GC-EAD, GC-MS, fruit volatile compounds

## Abstract

Phytophagous insects are capable of detecting and locating suitable hosts, which emit volatile compounds. Polyphagous species appear to have a complex olfactory strategy given that their numerous hosts have diverse emission profiles. In particular, their hosts’ volatile emissions share some of the same compounds, providing chemical bridges between them. However, the behavioural plasticity observed in insect host selection suggests that other volatiles have a complementary role. Here we explore how the specialization of polyphagous Tephritidae fruit fly in detecting and discriminating between host fruits has driven their chemical selectivity. The volatile emissions from intact or mechanically damaged fruit of 28 different species were analysed using gas chromatography-mass spectrometry and fed into a neuronal model of an olfactory system. We predicted *in silico* a functional trade-off between the two tasks, with optimal performance depending on detecting a higher proportion of shared fruit compounds, but with lower sensitivity compared to unshared compounds, or vice-versa. Using triple point electroantennography and a behavioural assay, we studied the olfactory response of Tephritidae fruit fly species that oviposit on fruit. Amplitude of the olfactory responses of eight species were negatively correlated with the compound’s degree of sharedness among fruit emissions, while response probability was previously shown to correlate positively with a similar metric. A dose-dependent switch in the fly’s preference confirmed the ecological importance of both shared and unshared fruit compounds. Thus, we propose that insect olfactory systems are chemically tuned to detect suitable hosts and accurately discriminate between them.

## Introduction

Phytophagous insects need to find a suitable host plant to fulfil their primary functions, such as feeding, mating and oviposition. They rely on their olfactory ability to perceive and identify their host by detecting its volatile compounds (Cardé and Willis, 2008; Schoonhoven et al., 2005). Their specialized peripheral olfactory system allows them to detect host compounds and integrate corresponding signals in their environment. However, in a natural setting there is a miasma of volatile compounds, which means that identifying a host is a complex task requiring an effective strategy. The volatile signature that an insect identifies as a host could correspond to particular ratios of a set of compounds or to individual compounds that are highly characteristic of the host (Bruce et al., 2005; Bruce and Pickett, 2011). Understanding the olfactory strategy involves deciphering both the chemical and sensory components of the communication system, which may facilitate the development of novel odour-based pest control strategies (e.g. Siderhurst and Jang, 2010).

Polyphagous species have a wide host range, which does not necessarily make their olfactory strategy more complex. It is possible that their preferred volatile signatures fit many plants, including numerous suitable hosts. Focusing on this hypothesis led to the discovery that Miridae or Tephritidae insect species are attracted to sets of the compounds, all of which are emitted by their many hosts (Biasazin et al., 2019; Cunningham, 2012a; Cunningham et al., 2016; Pan et al., 2015). However, this does not fully explain why polyphagous species can change their host preference depending on their past experience (Cooley et al., 1986; Cunningham et al., 2001, 1998). This behavioural plasticity suggests that polyphagous species can differentiate between their hosts, which can not rely on the ability to identify host-shared compounds. We have yet to resolve the relative contribution of compounds that are host-shared and non-host-shared (herein species-specific).

There is little experimental data available to address this issue. The family Tephritidae, a group of major pests, includes a substantial contingent of polyphagous species (Clarke, 2016; He et al., 2021a; Starkie et al., 2022). Some have an extremely wide host range, for example, more than 250 different host plants have been reported for the Oriental fruit fly *Bactrocera dorsalis* (Hassani et al., 2016; Moquet et al., 2021). Biasazin and colleagues showed that four polyphagous fruit fly species from this family detect preferentially and are strongly attracted to compounds that are shared between four different host fruits (Biasazin et al., 2019). As could be expected when dealing with highly polyphagous species, host selection plasticity has also been reported (Cooley et al., 1986; Cunningham et al., 2001; Manoukis et al., 2018; Segura et al., 2002).

The present study focuses on the evolutionary pressure on the peripheral olfactory system that led to the development of a polyphagous strategy in eight Tephritidae fruit fly species with wide and overlapping host ranges: *B. dorsalis*, *Bactrocera zonata*, *Dacus demmerezi*, *Ceratitis capitata*, *Ceratitis catoirii*, *Ceratitis quilicii*, *Neoceratitis cyanescens* and *Zeugodacus cucurbitae* (Charlery de la Masselière et al., 2017; Moquet et al., 2021). First, we explored headspaces from 28 highly divergent species of host fruit, by analysing the composition of their volatile emissions using gas chromatography coupled with mass spectrometry (GC-MS). Damaged fruit in orchards may emit new and potentially attractive compounds to fruit flies. Therefore, we sampled both intact fruit still on the tree and harvested fruit with mechanical damages. Using a semi-automated workflow for mass-spectrometry, we were able to isolate and identify several hundred compounds, for which we estimated the degree of sharedness among fruit emissions using alpha-diversity index. We then mapped antennal olfactory responses of females from the eight Tephritidae species to a set of synthetic fruit compounds. These were selected with a varying degree of alpha-diversity, using triple electroantennography (EAG3) (Ramiaranjatovo et al., 2023) and chopper-modulated GC, coupled with a triple electroantennogram detector (GC-EAD3). The resulting data allowed us to test whether the chemical tuning of the insect’s olfactory systems is correlated to the degree of compound sharedness among its hosts.

As a proxy for evolutionary pressure, we constructed a neuronal model of the peripheral olfactory system based on the likely composition of Tephritidae. We fed the fruit headspace data into the model and assessed how much the probability and the amplitude of antennal responses to species-specific and shared fruit compounds contribute to detecting and/or discriminating between host fruit species.

Lastly, the model’s prediction of a dose-dependent contribution of shared and species-specific fruit compounds led us to test the behavioural preference of female *B. dorsalis* using a dual choice trap, *ex situ*, with different doses of two sets of volatiles with a contrasting degree of alpha diversity.

## Results

### The volatile emissions of many fruit species partially overlap

To explore the diversity of the volatile emissions of 28 fruit species using ATD-GC-MS, we analysed both intact fruit (still on the tree) and sliced fruit (Figure 1). For intact fruit, we detected 511 compounds, of which 379 were identified and a further 55 were unidentified, but assigned to a chemical class. For the sliced fruit, we detected 665 compounds, of which 463 were identified and a further 38 were assigned to a chemical class. The main chemical classes were esters (13.31% and 18.35% of the total number of compounds for intact and sliced fruit, respectively) and terpenoids (14.87% and 13.38% of the total number of compounds for intact and sliced fruit, respectively). The remaining compounds were classified as: alcohol, aldehyde, amine, aromatic, carboxylic acid, ether, furanoid, green leaf volatile (GLV), hydrocarbon, ketone, organohalogen, organophosphorus, organosulfur and unknown (Figure 1E). The proportions of the different chemical families of compounds in intact and sliced fruit were significantly different (χ2 (15) = 31.084; p = 0.008). Table S1 and S2 provide the list of all the identified compounds.

**Figure 1.**
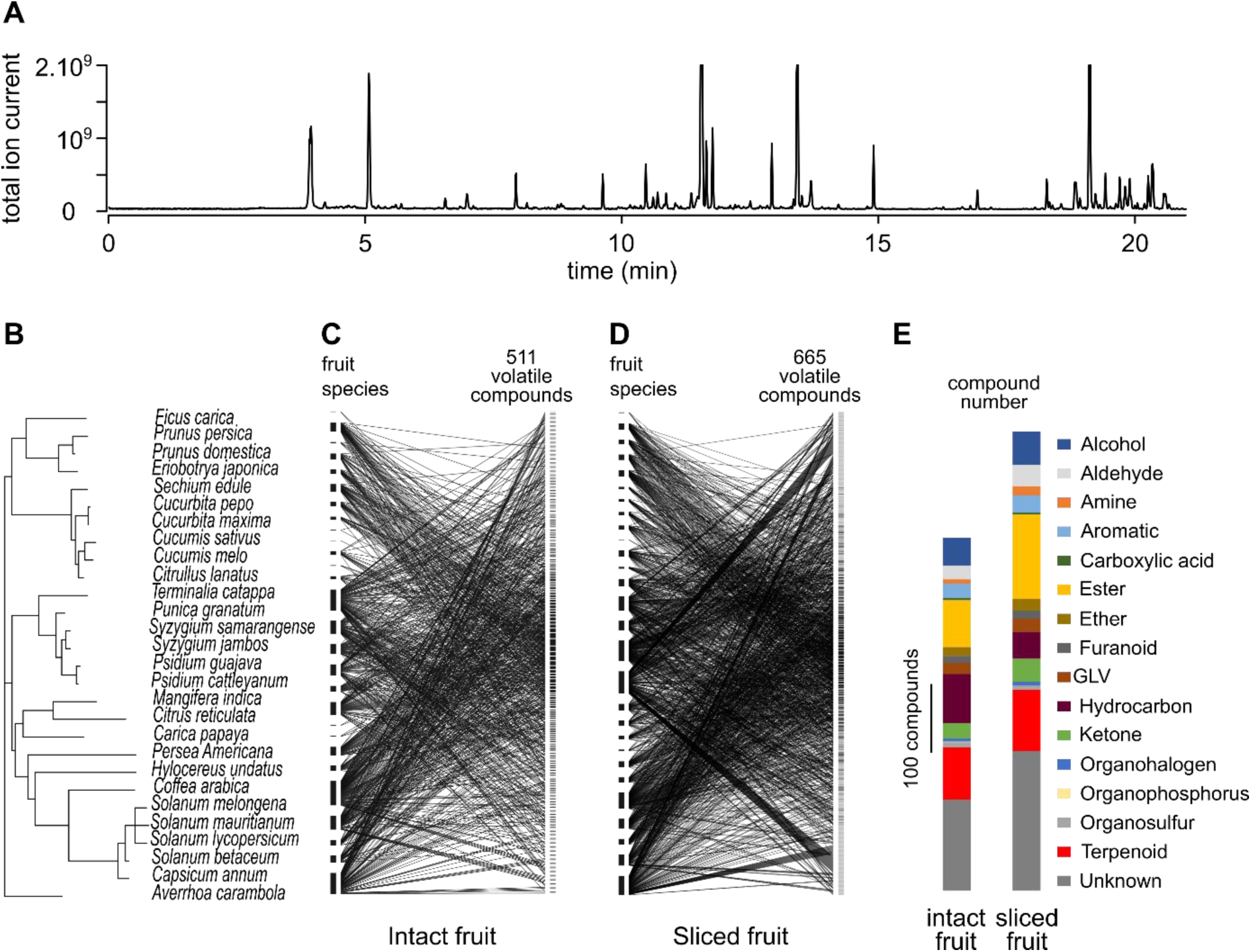
Diversity of volatile emissions produced by the 28 host fruit species of the Tephritidae studied. (A) Example of a chromatogram collected from the volatile emissions of a guava fruit (*Psidium guajava*). (B) Phylogeny of the 28 fruit species from which volatile emissions were collected. (C) Interaction network between the fruit species (left) and the 511 volatile compounds (right) collected from intact fruit. The fruit species are sorted in the same phylogenetic order as in panel (A), and the compounds are sorted by an index of fruit α-diversity, for which the maximal number is in the middle and the minimal (1 fruit) is at both extremities. (D) Interaction network between the fruit species (left) and the 665 volatile compounds (right) collected from sliced fruit, ordered as in panel (C). (E) Chemical classification of volatile compounds identified for the 28 fruit species. The two bars show the compounds of intact fruit (left) and sliced fruit (right), respectively. Each colour represents the chemical classes of compounds and the corresponding height is proportional to the number of compounds in each chemical class.

Despite the great diversity of fruit species included in this study, fruit emissions overlapped partially, with compounds emitted by 1 to 27 fruit species. We used indices of α-diversity among the volatile emissions of intact (α-div_IF_28) and sliced (α-div_SF_^28^) fruit to quantify the degree of sharedness. If there is a strong phylogenetic signal, closely related fruit species are likely to emit the same compounds. To test if high α-diversity values result from this type of phylogenetic autocorrelation, we computed the phylogenetic signal using the Abouheif’s *C_mean_* index for intact (C_mean, IF_^28^) and sliced (C_mean, SF_^28^) fruit samples. Indeed, we found a significant correlation between α-div_IF_28 and C_mean, IF_^28^ (F (1, 509) = 13.6; p < 0.001) and between α-div_SF_ ^28^ and C_mean, SF_ ^28^ (F (1, 663) = 19.3; p < 10^-4^). However, the variance explained by these models was low (r^2^ = 0.026 and r^2^ = 0.028, respectively) (Figure 2).

### Tephritidae olfactory responses are negatively correlated with the degree of sharedness of volatile compounds among fruit species

A set of 40 commercially available compounds, regularly distributed along the fruit α-diversity indices, was used to test the olfactory system of Tephritidae species (Table S3). We chose compounds from two chemical classes only, esters and terpenoids, given that we intended to explore the role of ecological rather than chemical variability. For these compounds, the indices of α-diversity, α-div_IF_28 and α-div_SF_ ^28^, were significantly correlated (F (1, 34) = 22.5; p<10^-4^). Moreover, α-div_SF_ ^28^ was not correlated with any of the compounds’ obvious chemical properties, namely boiling point (F (1, 38) = 2.1; p = 0.16), vapour pressure (F (1, 38) = 1.6; p = 0.21), lipophilicity estimated with log *Poct/wat* (ChemSketch, ACD/Labs) (F (1, 38) = 0.49; p = 0.49), or depression rate (Andersson et al., 2012) (F (1, 38) = 1.2; p = 0.28) (Figure S1).

**Figure 2.**
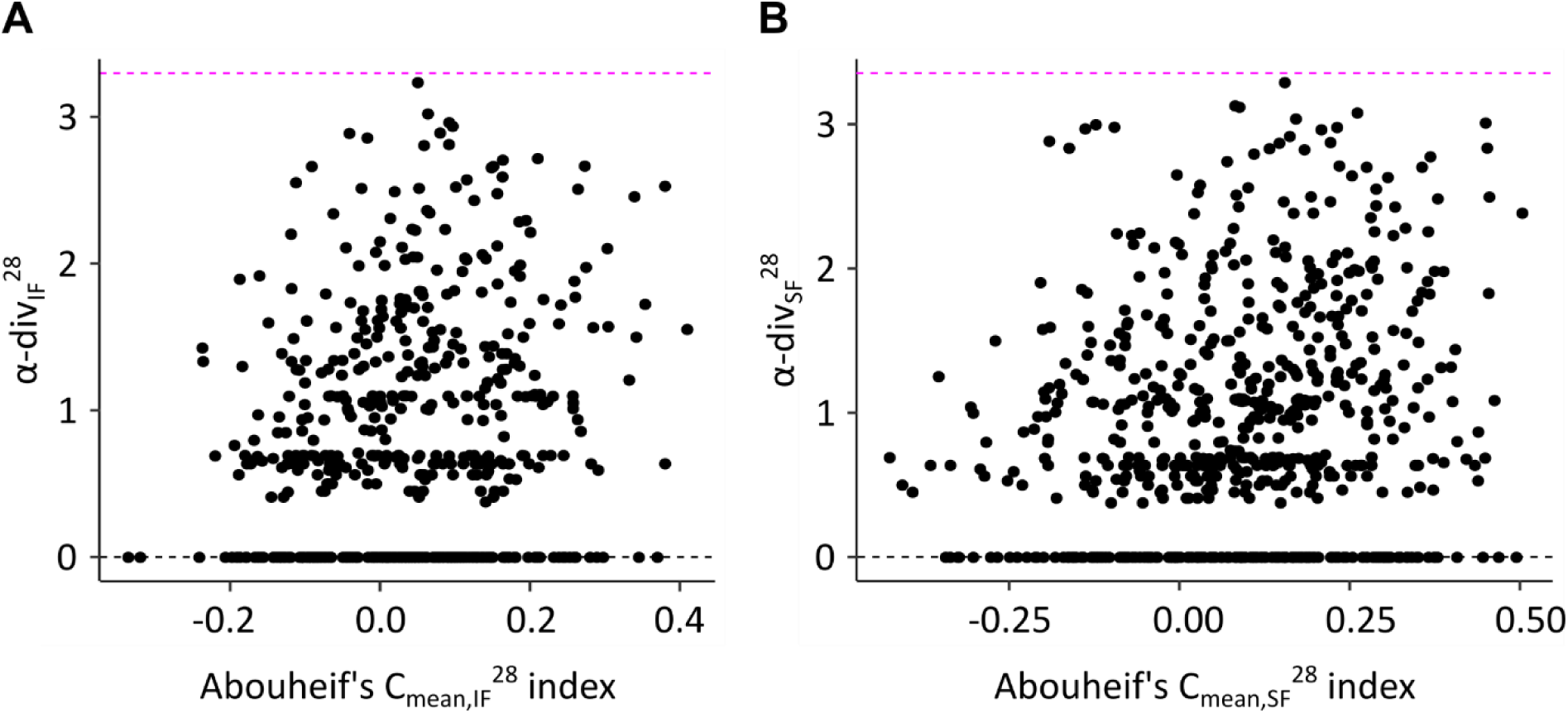
Relationship between α-diversity indices and phylogenetic signal Aubouheif’s C*mean* of fruit volatile compounds. Each point represents a compound for the emissions from intact (A) and sliced (B) fruit. Compounds with an α-diversity of 0 are emitted by only one fruit species. The magenta dashed line indicates the theoretical maximal α-diversity value reached if all fruit species emit the compound. A C*mean* of 0 means phylogenetic independence, while values above 0.2 are considered phylogenetically correlated.

The antennal responses to 30 of the compounds were measured in females from eight Tephritidae species with EAG3 at a dose of 10^-4^ (dilution v/v). To rule out possible experimental bias, we tested how this measure correlated with α-div_SF_ ^28^, using the compound depression rate as an independent variable. Antennal response was negatively correlated with α-divSF28 (F (1, 884) = 88.5; p < 10^-15^), with an interaction with species identity (F (7, 884) = 2.1; p =0.05). More precisely, it was significant in all species tested independently, except *B. zonata*, n = 4, F (1, 104) = 3.59; p = 0.06: *B. dorsalis*, n=4 individuals, F (1, 104) = 12.6; p < 0.001; *C. capitata*, n = 5, F (1, 130) = 24; p < 10^-5^; *C. catoirii*, n = 5, F (1, 130) = 39; p < 10^-8^; *C. quilicii*, n = 3, F (1, 78) = 13; p = 0.001; *D. demmerezi*, n = 4, F (1, 104) = 9; p = 0.003; *N. cyanescens*, n = 5, F (1, 130) = 15; p < 0.001; and *Z. cucurbitae*, n = 4, F (1, 104) = 9.8; p = 0.002 (Figure 3). Antennal response was also negatively correlated with α-div_IF_28 (F (1, 782) = 56; p <10^-13^), with an interaction with species identity (F (7, 782) = 2.9; p < 0.01). The effect was similar at a higher dose (10^-2^ dilution v/v) tested in *B. dorsalis*: the antennal response was still negatively correlated with α-div_SF_ ^28^ (n = 7 individuals, F (1, 182) = 23; p < 10^-5^) and with α-div_IF_ ^28^ (F (1, 161) = 17; p < 10^-4^, Figure S2).

**Figure 3.**
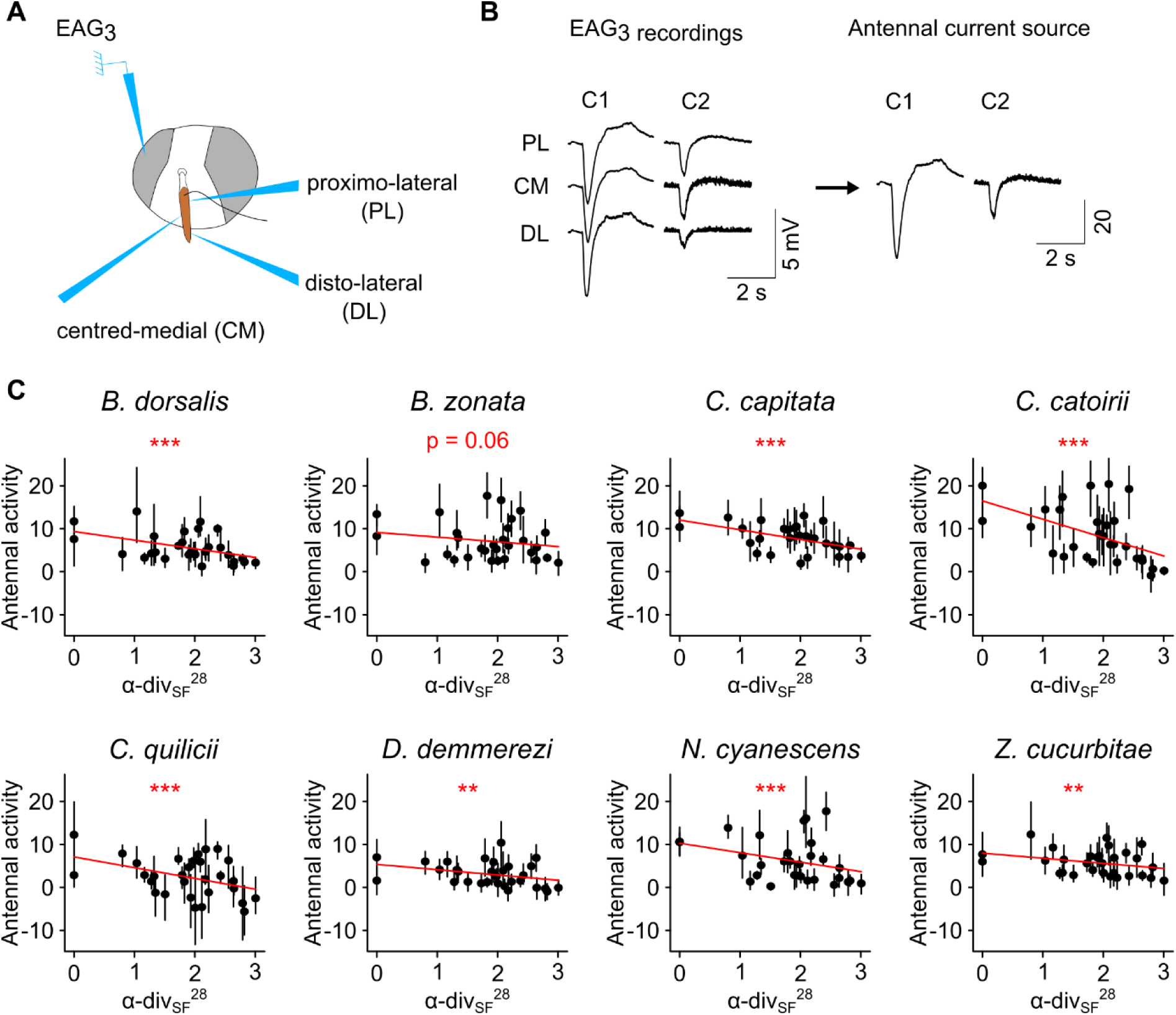
Antennal responses of eight Tephritidae species measured with EAG3 correlates with fruit α-diversity. (A) Schematic representation of the EAG3 mounting. (B) An example of EAG3 recording on individual female *B. dorsalis* stimulated with two compounds (C1 and C2). Left, the three signals represent the simultaneous recording at the three positions on the antenna. Right, the corresponding antennal current source are inferred from a CSD model and used to estimate the total antennal activity. (C) Antennal responses of females from eight Tephritidae species to synthetic compounds at dilution v/v: 10^-4^ depends on sliced fruit α-diversity. Points: mean value; bars: bootstrapped 95% c.i., n = 3 to 5 individuals per species. Red lines correspond to linear regression curves, whose significance is shown (** p<0.01; *** p<0.001).

The number of fruit species considered was of little consequence: we calculated indices of α-diversity among the volatile emissions of a subset of 13 fruit species (Table 1), reported to have a *B. dorsalis* infestation rate of above 5% in La Réunion (named α-div_IF_^13^ and α-div_SF_^13^ for intact and sliced fruit, respectively). We found a significant correlation between α-div_IF_^13^ and α-div_IF_28 (F (1, 34) = 79.8; p < 10^-9^) and between α-div_SF_ ^13^ and α-div_SF_ ^28^ (F (1, 38) = 128.4; p < 10^-13^, Figure S3). Again, antennal response was negatively correlated with α-div_SF_ ^13^ (F (1, 884) = 100; p <10^-15^), with an interaction with species identity (F (7, 884) = 2.1; p =0.05). It was significant in each species tested individually: *B. dorsalis*, F (1, 104) = 13.7; p < 0.001; *B. zonata*, F (1, 104) = 5.1; p = 0.03; *C. capitata*, F (1, 130) = 20; p < 10^-4^; *C. catoirii*, n = 5, F (1, 130) = 28; p < 10^-7^; *C. quilicii*, F (1, 78) = 19; p < 10^-4^; *D. demmerezi*, F (1, 104) = 14; p < 0.001; *N. cyanescens*, F (1, 130) = 18; p < 10^-4^; and *Z. cucurbitae*, n = 4, F (1, 104) = 19; p <10^-4^. Antennal response was also negatively correlated with α-div_IF_^13^ (F (1, 782) = 55; p <10^-12^), with an interaction with species identity (F (7, 884) = 2.0; p = 0.05). We then explored the EAD3 antennal response to 37 compounds. Here, the stimulations were provided through chopping modulated GC. This protocol induced more experimental bias, since antennal responses to GC-stimulations were strongly correlated with compound depletion rate (F (1, 280) = 29; p < 10^-6^), which might result from the chopping modulation. However, we found that the GC-EAD3 antennal response of the eight species was negatively correlated with div ^28^ (F (1, 264) = 10.5; p=0.001) and with div ^28^ (F (1, 232) = 30.1; p<10^-7^), with no interaction with species identity (F (7, 264) = 0.4; p=0.9 and F (7, 264) = 0.9; p=0.5, respectively). For each species tested independently, antennal response depended on a joint effect between div_IF_28 and compound depletion rate (Figure S4): *B. dorsalis*, n=4 individuals, F (1, 29) = 8.2; p < 0.01; but *B. zonata*, n = 4, F (1, 29) = 4.1; p = 0.05; *C. capitata*, n = 3, F (1, 29) = 11.8; p = 0.002; *C. catoirii*, n = 3, F (1, 29) = 4.8; p = 0.04; *C. quilicii*, n = 3, F (1, 29) = 9.3; p = 0.005; *D. demmerezi*, n = 3, F (1, 29) = 7.8; p = 0.01; *N. cyanescens*, n = 3, F (1, 29) = 9.2; p =0.005; and *Z. cucurbitae*, n = 4, F (1, 29) = 24.4; p <10^-4^. The effect was similar at a higher dose (1 μg) tested in *B. dorsalis*: antennal response depended on a joint effect between div_IF_28 and compound depletion rate (n = 5 individuals, F (1, 29) = 8.2; p <0.01). To sum up with regard to GC-EAD3 data, the main effect was a negative correlation between antennal response and div_IF_^13^ for compounds with a low depletion rate.

### A neuronal model predicts a contrasted use of species-specific and shared fruit compounds for odour-based fruit detection and discrimination

The amplitude of EAG responses, as observed in this study, and the probability that a compound induces an EAG response, as reported previously (Biasazin et al., 2019), appear to be inversely correlated with compound sharedness among fruit. We hypothesised that the bidirectional property of olfactory tuning in these species was selected both by the ecological need to detect a large number of hosts, encouraging responses to host-shared compounds, and the ecological need to discriminate host species, encouraging responses to species-specific compounds. Neuronal modelling was used to test this. We built 120,000 random neuronal models of a peripheral olfactory system (Figure 4A). The GC-MS data for intact or sliced fruit samples were used as input odours for these models in order to consider the natural variability of fruit emissions within and between fruit species, and the natural distribution of the degree of sharedness of volatile compounds in fruit emissions. For each model, we estimated the efficiency of the olfactory system to detect and discriminate fruit using specially designed indices (Figure 4B). The performance of the models was heterogeneous, allowing us to explore the typical chemical tuning of the most efficient olfactory systems for either detection, discrimination or both. Not surprisingly, models with 20 ORs discriminated fruit better than models with 10 ORs (F (1, df > 10^5^) = 258,197; p<10^-15^), and the fruit detectability index was correlated with the number of compounds detected per OR (F (2, df > 10^5^) = 921,033; p < 10^-15^). For cross-comparison, we standardized the distributions (expressed as z-scores) for each combination of number of ORs, number of compounds detected per OR and input data sets (intact or sliced fruit emissions).

**Figure 4.**
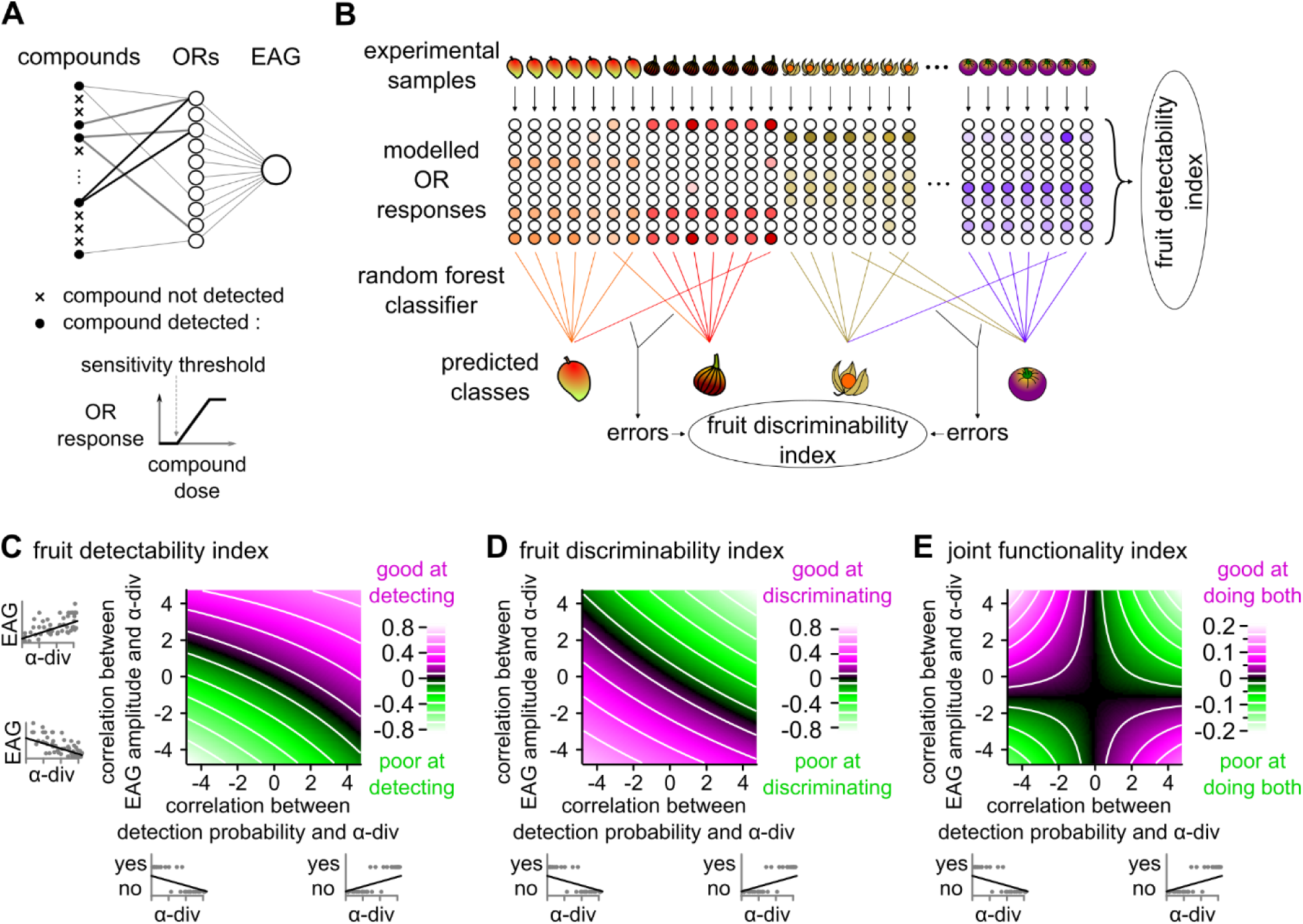
Neuronal model reveals a functional trade-off between detection of and discrimination between fruit species. (A) Schematic representation of the parameters included in one random model of an insect’s olfactory system. Each olfactory receptor (OR) is sensitive to a random set of compounds, as shown by a simplified dose-response curve, whose threshold and dynamic range have been drawn randomly. (B) Schematic depiction of the algorithms used to calculate the fruit detectability index and the fruit discriminability index for one random model. For this, experimental data corresponding to intact or sliced fruit samples were used as an input (top) and the modelled OR responses were calculated (middle). The fruit detectability index was the sum of all OR responses to all volatile samples (right). The fruit-species discriminability index (bottom) was calculated from the proportion of fruit species misclassified by a random forest algorithm applied on the OR response patterns. (C) Fruit detectability index (color code) is correlated both with the correlation coefficient between EAG amplitude and fruit α-diversity and the correlation cefficient between the probability to detect a compound and fruit α-diversity (linear model p<10^-15^ in both cases, all variables are expressed in z-score units). (D) Fruit discriminability index (color code) is correlated both with the correlation coefficient between EAG amplitude and fruit α-diversity and the correlation cefficient between the probability to detect a compound and fruit α-diversity (linear model p<10^-15^ in both cases, all variables are expressed in z-score units). (E) For models to efficiently detect and discriminate fruit at the same time, amplitude and probability of response should be inversely correlated with fruit α-diversity (color code: joint functionality index, all variables are expressed in z-score).

**Table 1:** List of the 28 fruit species studied. Asterisks * indicate the 13 species reported to have a *B. dorsalis* infestation rate of over 5%.

| Fruit families | Fruit species | Common names |
| --- | --- | --- |
| Anacardiaceae | *Mangifera indica* * | Mango |
| Cactaceae | *Hylocereus undatus* * | Dragon fruit |
| Caricaceae | *Carica papaya* * | Papaya |
| Combretaceae | *Terminalia catappa* * | Indian almond |
| Cucurbitaceae | *Sechium edule* | Chayote |
| Cucurbitaceae | *Cucurbita maxima* | Pumpkin |
| Cucurbitaceae | *Cucumis sativus* | Cucumber |
| Cucurbitaceae | *Cucurbita pepo* | Zucchini |
| Cucurbitaceae | *Cucumis melo* | Melon |
| Cucurbitaceae | *Citrullus lanatus* | Watermelon |
| Lauraceae | *Persea americana* * | Avocado |
| Lythraceae | *Punica granatum* | Pomegranate |
| Moraceae | *Ficus carica* * | Fig |
| Myrtaceae | *Psidium guajava* * | Guava |
| Myrtaceae | *Psidium cattleianum* * | Strawberry guava |
| Myrtaceae | *Syzygium samarangense* * | Java apple |
| Myrtaceae | *Syzygium jambos* * | Rose apple |
| Oxalidaceae | *Averrhoa carambola* | Star fruit |
| Rosaceae | *Eriobotrya japonica* * | Loquat |
| Rosaceae | *Prunus persica* * | Peach |
| Rosaceae | *Prunus domestica* * | Plum |
| Rubiaceae | *Coffea arabica* | Coffee |
| Rutaceae | *Citrus reticulata* | Tangor |
| Solanaceae | *Solanum melongena* | Aubergine/Eggplant |
| Solanaceae | *Solanum mauritianum* | Bugweed |
| Solanaceae | *Capsicum annuum* | Chilli |
| Solanaceae | *Solanum betaceum* | Tree tomato |
| Solanaceae | *Solanum lycopersicum* | Tomato |

We calculated, for each model, the correlation coefficient between the probability for a compound to be detected and fruit α-diversity, and the correlation coefficient between the amplitude of the EAG response to those detected compounds, at four different doses, and fruit α-diversity. By design, the two variables were independent (F (1, df > 10^5^) = 0.02; p = 0.89). The fruit detectability index depended significantly on the two correlation coefficients (Figure 4D, F (1, df > 10^5^) = 1,238 and 3,939; p < 10^-15^ and 10^-15^, respectively) with a significant interaction (F (1, df > 10^5^) = 11; p < 0.001). The fruit discriminability index also depended significantly on the two correlation coefficients (Figure 4D, F (1, df > 10^5^) = 1,988 and 4,020; p < 10^-15^ and 10^-15^, respectively) with a significant interaction (F (1, df > 105) = 5.6; p =0.018). More specifically, the two response parameters (probability and amplitude) were positively correlated with fruit α-diversity in the most efficient models at detecting fruits, and negatively correlated with it in the most efficient models at discriminating fruits. This functional trade-off was robust to the model parameters, as it was observed for any OR numbers, numbers of detected compounds per OR and intact or sliced fruit input data, or stimulation dose tested (Figure S5). The effect on EAG amplitude depended mostly on the sensitivity thresholds to detected compounds (Figure S6) and, to a lesser extent, the dynamic range of detected compounds.

We then calculated an index of joint functionality, assessing how much the olfactory system is efficient at doing both functions simultaneously. This index depended on a joint effect between both correlation coefficients (Figure 4F, F (1, df > 10^5^) = 21; p < 10^-5^). This effect was robust to the model parameters, even though it was weaker for EAG responses to low doses of compound, or for those models with a high proportion of detected compounds (Figure S5). Specifically, it points to two possibilities for an olfactory system to be efficient at detecting and discriminating fruit at the same time: probability of detecting compounds and response amplitude should be inversely correlated, in a direction or the other, with fruit α-diversity.

### Female *B. dorsalis* display a dose-dependent switch in behavioural preference between shared and species-specific fruit compounds

Overall, the olfactory tuning of Tephritidae antennae is consistent with the hypothesis that the ability to discriminate fruit species has an ecological role, as predicted by the models. A matching prediction would be that at low doses, flies would only perceive species-specific fruit compounds. We designed a behavioural assay involving a dual choice *ex situ* trap to test this (Figure 5A). We selected five species-specific fruit compounds and five shared compounds, all of which were small esters to ensure low chemical variability (Table 2). A behavioural experiment showed that female *B. dorsalis* were more attracted to a blend of the 10 compounds, whose emission rates were homogenized, than to a negative control (GLMM: χ2 (1, N = 11) = 28.845; p < 0.001; mean proportion of flies trapped after 125 min: 0.185 (95% CI 0.118; 0.252) and 0.003 (95% CI -0.003; 0.009), respectively; Figure 5B). Then, we tested the comparative attractiveness of the two sub-blends of five species-specific fruit compounds and five shared compounds. At the lowest dose tested (dilution v/v: 10^-4^), insects were more attracted to the species-specific compound blend than to the shared compound blend (GLMM: χ2 (1, N = 46) = 214.617; p < 0.001; mean proportion of flies trapped after 240 min: 0.186 (95% CI 0.118; 0.254) and 0.048 (95% CI 0.027; 0.070), respectively; Figure 5C). At a higher dose (dilution v/v: 10^-3^), a reversal of the trend was observed, i.e. females preferred the shared compound blend over the species-specific compound blend (GLMM: χ2 (1, N = 60) = 38.355; p < 0.001; mean proportion of flies trapped after 240 min: 0.133 (95% CI 0.098; 0.168) and 0.109 (95% CI 0.076; 0.142), respectively; Figure 5D).

**Figure 5.**
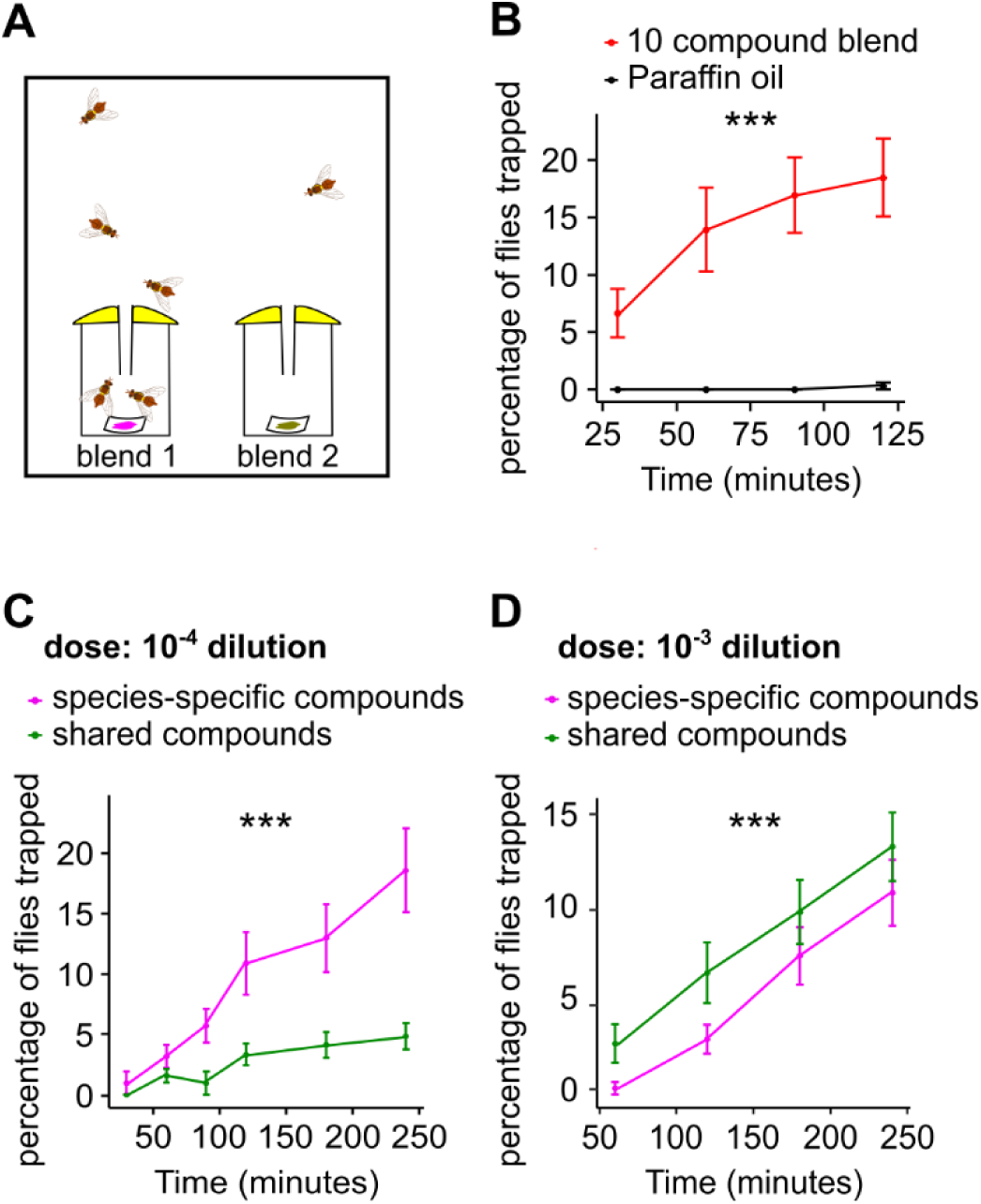
*B. dorsalis* behavioural preference between species-specific and shared fruit compounds is dose-dependent. (A) Schematic drawing of the *ex situ* trapping bioassay. (B) Dual choice tests between a blend of 10 compounds diluted at 10^-4^ v/v in mineral oil (red) and mineral oil (black); values are mean +- SE. ***: significant difference between the two blends GLMM, p < 0.001. (C) Dual choice tests between a blend of five shared compounds (in green) and a blend of five specific compounds (in magenta) tested at the dilution v/v: 10^-4^. Same convention as in panel (B). (D) Dual choice tests between the same blends as in panel (C) diluted at 10^-3^ v/v. Same conventions as in panel (B).

**Table 2:** List of compounds used for behavioural tests. The last column shows a list of fruit species in which the compound was found in sliced fruit samples only and not in intact fruit.

| blend | CAS | compound | Found in the emissions of: |  |
| --- | --- | --- | --- | --- |
|  |  |  | intact fruits | sliced fruit only |
| shared compounds | 3681-71-8 | cis-3-hexenyl acetate | *Indian almond, strawberry guava, peach, plum* | *bugweed, coffee, loquat, mango, melon, star fruit, tomato* |
|  | 141-78-6 | ethyl acetate | *bugweed, coffee, Indian almond, loquat, mango, peach, strawberry guava, tree tomato* | *chilli, guava, melon, rose apple, star fruit, tomato* |
|  | 105-54-4 | ethyl butyrate | *strawberry guava, rose apple, mango* | *bugweed, melon, peach, plum, star fruit, tree tomato* |
|  | 105-37-3 | ethyl propionate | *bugweed, rose apple, tomato, papaya* | *guava, melon, plum, star fruit* |
|  | 142-92-7 | hexyl acetate | *chilli, eggplant, Indian almond, papaya, peach, pumpkin, star fruit, strawberry guava* | *plum* |
| species-specific compounds | 109-21-7 | butyl butyrate | *Plum* | - |
|  | 868-57-5 | methyl 2-methylbutyrate | *rose apple* | *melon, star fruit* |
|  | 106-73-0 | methyl heptanoate | *star fruit* | - |
|  | 111-11-5 | methyl octanoate | *peach, star fruit* | *guava, strawberry guava* |
|  | 1191-16-8 | prenyl acetate | *Indian almond* | *guava, strawberry guava* |

## Discussion

In this paper we used chemical analysis and modelling, combined with electrophysiological and behavioural approaches to explore the chemical interactions between Tephritidae species and 28 of their host fruit species. By analysing the chemical composition of volatile emissions from 28 Tephritidae host fruit, either intact (on the tree) or after mechanical damage, we identified a considerable number of compounds. As expected, damaged fruit produced more volatile compounds than intact fruit (Beck et al., 2008; Geervliet et al., 1994; Meiners, 2015). The volatile emissions were dominated by terpenoids and esters, which are characteristic of volatile fruit emissions and are easily detected by insect olfactory receptors (Dweck et al., 2018; El Hadi et al., 2013). Although each fruit species has its own chemical profile, there is considerable overlap between species. The degree of sharedness among fruit species (fruit α-diversity) varied, some compounds were specific to a single species, while others were emitted by most of the fruit species considered. We found a significant negative correlation between the antennal responses of several Tephritidae species and volatile α-diversity. The effect was robust, regardless of how the α-diversity index was calculated by considering: all 28 fruit species or a subset of 13 species, with a high infestation rate of generalist species; and intact or mechanically damaged fruit. Lastly, we found that mature female *Bactrocera dorsalis* demonstrated a dose-dependent behavioural preference when we compared a blend of species-specific fruit compounds and a blend of shared compounds. Below, we will discuss how our observations and neuronal modelling support the alternative hypothesis regarding the olfactory strategy used by Tephritidae to explore their wide host range.

The main hypothesis suggests that since Tephritidae females have an ecological need to detect a large number of suitable hosts for oviposition, their olfactory system should be tuned for compounds shared by different fruit species. This type of behaviour would allow females to identify a wide host range with minimum effort in terms of information processing and, therefore, minimum energy consumption (Bernays, 2001; Cunningham, 2012b; Niven and Laughlin, 2008). Supporting this theory, the addition of three compounds identified in ripe guava considerably increased *Bactrocera tryoni*’s attraction to several non-preferred hosts (Cunningham et al., 2016). The authors suggest that these compounds could be a generic signature for ripe fruit. Furthermore, a blend of volatile compounds shared between four phylogenetically distant fruit species was attractive for *Bactrocera dorsalis* and *Zeugodacus cucurbitae* (Biasazin et al., 2019). The hypothesis raises a number of issues, which we tested. First, do phylogenetically distant fruit species emit the same volatile compounds? Clarke predicted a substantial proportion of shared compounds was an adaptation to generalist seed dispersers (Clarke, 2017). We found that fruit α-diversity was only weakly correlated with fruit phylogeny, i.e. a number of identical compounds were emitted by fruit that did not belong to the same clade. Relying on shared compounds could explain why tephritid host associations do not correlate with fruit phylogeny (Charlery de la Masselière et al., 2017). Second, do generalist species prefer shared compounds over species-specific fruit compounds? Our observations reveal a preference for shared compounds in *B. dorsalis*, but only for the highest dose tested. Third, do shared compounds induce a stronger antennal response and, therefore, are they overrepresented compared to species-specific compounds? Our neuronal model confirms that this should be the case and suggests two testable predictions: the proportion of compounds detected by the olfactory systems should correlate positively with the compound’s degree of sharedness among fruit species, which was indeed observed in another study on Tephritidae (Biasazin et al., 2019); the response amplitude to detected compounds should also correlate positively with the compound’s degree of sharedness among fruit species, but here we observed a negative correlation. Our observation is unlikely to be an experimental artefact because a bias due to the heterogeneous volatility of the tested compounds should not make a difference. Indeed, we found no correlation between fruit α-diversity and boiling point, lipophilicity or depletion rate (Andersson et al., 2012). In addition, the effect was observed both with a passive and GC-induced vaporization of compounds. The use of triple electroantennography reduces the risk of uneven sampling of the olfactory receptor^23,24^. Artefactual responses to impurities have been reported (Paoli et al., 2017; Schorkopf et al., 2019), but were refuted here by the GC-EAD analysis of the same compounds.

When faced with various volatile compound emitters, insects must be able to identify their hosts’ volatile compounds from a complex environment, which may include volatile emissions from non-host plants (Bruce et al., 2005; Bruce and Pickett, 2011). Their olfactory systems may be tuned to host-specific compounds. Some compounds shared between fruit species may also be emitted by other plant parts and non-host species. Therefore, they may be widely distributed in the natural environment where the species evolved and, thus, be neglected by the insect. Cis-3-hexenyl acetate is a case in point. Although emitted by many fruit species, this compound is also emitted by leaves in response to herbivore damage (Scala et al., 2013). While plant compounds like this may have high fruit α-diversity, they may have little ecological relevance for insects, if any. In this scenario, a strong antennal response would only be expected for shared compounds that are specifically emitted by fruit. However, the hypothesis would have predicted a low response to shared plant compounds and species-specific host compounds rather than the negative correlation that we observed. Moreover, shared compounds that are specific to fruit should be attractive to insects, irrespective of the dose. We observed a preference to species-specific fruit compounds at low doses, which suggests that another factor may be involved.

Our results led us to consider the alternative hypothesis, namely that polyphagous insects have an ecological need to discriminate between host species. The ability to change host preference to adapt to a dynamic environmental context provides evolutionary advantages, which have been widely discussed (Cunningham et al., 2001; Cunningham and West, 2008; Jaenike, 1978). Host preference in Tephritidae can change, as demonstrated by the comparative analysis of wild and laboratory-reared individuals of the medfly species, *Ceratitis capitata* (Joachim-Bravo et al., 2001). A switch in host preference can be induced in *C. capitata* females (Cooley et al., 1986) and adult attraction behaviour can be conditioned (Liu et al., 2018) or influenced by larval diet (Manoukis et al., 2018) in *B. dorsalis*. As an adaptive consequence of learning, host preference of *C. capitata* changes depending on host abundance (Segura et al., 2002), as predicted by Jaenike’s optimal oviposition behaviour theory (Jaenike, 1978). A prerequisite to host preference plasticity is the ability to discriminate hosts. We challenged the hypothesis that Tephritidae rely on species-specific fruit compounds to discriminate fruit species using a neuronal model of olfactory detection. The model suggested that ecological pressure to discriminate between fruit species would result in lower sensitivity thresholds for species-specific compounds compared to shared compounds. This phenomenon would generate a negative correlation between antennal response and fruit α-diversity. It is important to note that while our modelling formally validates a hypothesis that is consistent with our experimental observations, it does not constitute proof *per se*.

Given the functional trade-off suggested by the model, which olfactory tuning would be selected by the most likely hypothesis that species have an ecological need both to detect a diversity of hosts and to discriminate between them? The modelling suggests a possibility: the detection of a higher proportion of shared compounds than species-specific compounds, but with lower olfactory sensitivity. Thus, our observations add an additional level of complexity to the olfactory strategies of polyphagous species. Moreover, the low antennal response to shared compounds does not mean that insects neglect them, but that high sensitivity is not required to detect them given that they are frequently encountered in the insects’ natural environment. Therefore, we expected a dose-dependent switch in preference, which is exactly what we observed in our behavioural assay. It is important to note that blends of species-specific fruit compounds (i.e. with no shared compounds) do not exist in nature. Hence, we consider the effect as a sensory illusion, which stems from an ability to discriminate between hosts. To conclude, we postulate that insects detect hosts through host-shared compounds and choose between hosts through species-specific compounds. More generally, we expect that the three functions, namely, host detection, host vs non-host selection and discrimination between hosts help shape the chemical tuning of insects’ olfactory systems.

Surprisingly, we found a negative correlation between antennal response and fruit α-diversity in five polyphagous species and three oligophagous species of the tribes, Dacini and Ceratitidini. This suggests that the trait is conserved in Tephritidae, regardless of their ecology. This echoes the reported attraction of *Z. cucurbitae* and *B. dorsalis* to a blend of compounds shared by different fruit species (Biasazin et al., 2019). Two hypotheses can explain this. First, the oligophagous species that we studied may not be truly oligophagous. The Cucurbitaceae specialist, *Z. cucurbitae*, is sometimes considered to be polyphagous (Starkie et al., 2022), following reports of infestations in several fruit families including, Anacardiaceae, Caricaceae, Combretaceae, Moraceae, Solanaceae, etc. (Charlery de la Masselière et al., 2017; Dhillon et al., 2005; Kambura et al., 2018; Mcquate et al., 2015; Vayssières et al., 2007). Infestations of other fruit families were also reported for Cucurbitaceae specialists from the Dacus genus (Kambura et al., 2018; Vayssières et al., 2007) and for the Solanaceae specialist, *N. cyanescens* (Charlery de la Masselière et al., 2017). Secondly, phylogenetic analysis of host range diversity within the Dacini tribe revealed multiple episodes of host range switch, which involved broadening or reducing host range (He et al., 2021b; Starkie et al., 2022; Virgilio et al., 2009). Since peripheral olfactory systems are generally conserved across Tephritidae (Jacob et al., 2017), oligophagous species might have a polyphagous-type olfactory system due to phylogenetic inertia. To conclude, if the olfactory strategies we observed are related to polyphagy, similar analyses should be conducted on truly monophagous species or on polyphagous frugivores from other insect orders.

Understanding the olfactory strategies adopted by phytophagous insects that allow them to recognize their host plant not only provides fundamental insights into ecology and evolution, but it will also facilitate the effective design of attractants for pest control (Agelopoulos et al., 1999). When it comes to efficient trapping, our data suggest that tephritids may not have a high sensitivity to a blend of compounds shared by different fruit species. Inversely, a blend of species-specific fruit compounds may not be sufficiently attractive compared to a mixed blend of compounds emitted naturally by various host fruit species. Thus, it would be interesting to take our study further, for example, by combining compounds shared by different fruit species with species-specific fruit compounds chosen from the most attractive fruit. This type of blend would be attractive at both high and low doses and, by extension, at any distance from the lure. Identifying the optimum ratio of compounds in the blend is also a challenge, given the variation between fruit emissions. The ratio of shared compounds in a blend could either imitate an attractive fruit or correspond to the ratio of compounds most shared by host fruit species.

In conclusion, this study considers two non-exclusive hypotheses regarding the olfactory strategies developed by fruit flies, which allow them to recognize their plant hosts. Our study supports the view that tephritids’ olfactory systems are not only designed to detect hosts, but also to discriminate between them, which means they are efficient and flexible in their search for hosts.

## Materials and methods

### Insects

Female individuals of eight species of Tephritidae were used for the electrophysiological experiments: *Bactrocera dorsalis* (oriental fruit fly), *Bactrocera zonata* (peach fruit fly), *Ceratitis capitata* (medfly), *Ceratitis catoirii* (Mascarene fruit fly), *Ceratitis quilicii* (Cape fruit fly), *Dacus demmerezi* (Indian Ocean cucumber fruit fly), *Neoceratitis cyanescens* (tomato fruit fly) and *Zeugodacus cucurbitae* (melon fly). They originated from laboratory strains, which were reared according to previously published procedures (Charlery de la Masselière et al., 2017; Hafsi et al., 2016). The individuals selected for electrophysiology had lived for respectively 12-15, 28-32, 10-14, 21-26, 13-17, 11-15, 10-16 and 25-31 days after emergence and belonged to the following generations: 62-64^th^, 179-180^th^, 107-108^th^, 311-312^th^, 77-79^th^, 59^th^, 64-65^th^ and 7^th^.

The behavioural experiments were conducted on female individuals of *B. dorsalis* from the 76-78^th^ generations, 18-25 days after emergence.

### Plant materials and collection of volatile compounds

Fruit volatile compounds were collected using dynamic headspace extraction. We put individual or clusters of fruit into nalophan sample bags (SenseTrading, Gröningen, Netherlands) that were 47 cm wide. The bags were sealed with plastic ties at both ends to create a headspace. The air content of the headspace was sucked out at 100 mL.min^-1^ for two hours using an adsorption tube filled with Tenax GR (Supelco, Bellefonte, USA) and a portable membrane pump (Spectrex PAS-500; Spectrex, Redwood City, CA, USA). For each sample, a negative control sample was performed with an empty nalophan bag.

In this study, we selected 28 fruit species considered to be potential hosts of Tephritidae (Moquet et al., 2021) (Table 1). Headspace samples were extracted from intact fruit on the tree or from sliced fruit at a ripening stage suitable for oviposition. For each fruit species, sliced or unsliced, 3 to 8 (median 7) replicates were performed, with a total of 193 samples for intact fruit and 176 for sliced fruit.

### Gas chromatography

Tenax traps were stored at room temperature and analysed within seven days using an automated thermal desorber (ATD, Turbomatrix 350, Perkin Elmer, USA), coupled with GC-MS (Clarus 580 GC and SQ8T MS, Perkin-Elmer, USA). The parameters of the ATD were as follows: primary desorption 5 min at 250°C, flow 50 mL.min-1; no inlet split; up to an Air Monitoring trap kept at 2°C; secondary desorption 1 mL.min-1 for 5 min, and a constant flow of 3 mL.min-1. An Elite-5MS capillary column (60 m x 0.25 mm i.d., 0.25 µm film thickness, Perkin-Elmer, USA), was used for chromatography. The oven was programmed from 50 to 270°C at 8°C.min-1. Helium was used as a carrier gas with a constant flow of 0.21 mL.min-1.

### Triple electroantennography EAG3 bioassays

Electrophysiology experiments were conducted using the EAG3 technique (Ramiaranjatovo et al., 2023) (Figure 4A). EAG3 improves the estimation of the insect’s olfactory responses by simultaneously recording at three antennal positions. A live fly was immobilized in a 1,000 µL pipette tip, with its head exposed. The base of the head was fixed with dental wax (Enta Periphery Wax, Netherlands). For species of the Dacini tribe with elongated antennae, the first two antennal segments, which do not have olfactory functions, were stabilized with a silicone adhesive elastomer “Kwik-Sil” (World Precision Instruments, Inc., Sarasota, USA). Four electrodes, consisting of a chloride silver wire inserted into a glass micropipette (tip diameter 1-2 μm), filled with an electrolyte solution (120 mM NaCl, 5 mM KCl, 1 mM CaCl2, 4 mM MgCl2, and 10 mM HEPES), were used to perform the EAG3 recording. The recording electrodes were placed in the latero-proximal, medio-central and latero-distal regions of the right funiculus. The reference electrode was inserted in the middle of the left eye. The recording electrodes were connected to a four-channel differential AC amplifier (Model 1,700, A-M system, Carlsborg, USA) which amplified (gain x 1,000) and filtered the signals between 1 and 1,000 Hz. Data acquisition was fitted by digitalizing the signals at 500 Hz with an acquisition card (NI 9215, National instruments, France), associated with Labview software (National instruments, France). Data were analysed using a current source density model to infer the total antennal response from the EAG signal recorded at three positions (Jacob, 2018; Ramiaranjatovo et al., 2023).

Stimuli were delivered through a glass chamber (20 cm x 5 mm i.dFID.). The tube had a small hole 5 cm upstream its outlet into which the odour source was inserted. A continuous humidified and charcoal-filtered airflow (20 mL.s^-1^) across the glass chamber reached the insect, which was 1 cm from the tube outlet. The insect was stimulated using an odour cartridge consisting of a Pasteur pipette with a square of filter paper loaded with 1 µL of a solution of a synthetic compound diluted at 10^-4^ v/v or 10^-2^ v/v in mineral oil. The Pasteur pipette was inserted into the small hole in the glass chamber and exposed to a puff of air for 200 ms (3 mL.s^-1^), that was regulated with an electro-valve (LHDA-1233215-H, Lee Company, France), and controlled by a digital module (NI 9472, National Instruments, Nanterre, France), and Labview software (National Instruments, France). The experiment consisted of passing 30 compounds (listed in Table S3) individually in a random order with a time interval of 1 minute between each stimulation. A negative control stimulation (mineral oil) and a positive stimulation (1-octen-3-ol) were performed before and after each experiment.

### Coupling chopped gas chromatograph - triple electroantennography detector (GC-EAD3)

For GC-EAD3 experiments, EAG3 antennal preparation was used as a detector in parallel with the flame ionization detector (FID). A fused silica Restek Rxi-5 ms column (30 m x 0.32 mm i.d., 0.25 µm film thickness, Restek, Lisses, France) was used and heated as follows: 50°C for 1 min, then 6.5°C.min^-1^ to 200°C and 40°C.min^-1^ to 250°C, held for 5 min. The temperature of the FID was set to 250°C. Helium N60 was used as the carrier gas. To improve the signal to noise ratio, the GC system was equipped with a D-Swafer Dean’s switch (PerkinElmer, Inc., USA) at the column output, as described previously (Ramiaranjatovo et al., 2023). The device allows for the GC signal to be chopped with the alternate emission of the effluent to both the EAG3 and FID detectors (Myrick and Baker, 2018; Ramiaranjatovo et al., 2023). The chop frequency was regulated to 1 Hz with Labview software (National Instruments, France) and a digital module (NI 9472, National Instruments, Nanterre, France). The signal was demodulated at the chopping frequency (1 Hz) before a similar analysis as for the EAG3 recordings (Ramiaranjatovo et al., 2023). The sample was fed into the glass chamber using a 3 m long heated transfer line (Antelia, Dardilly, France). To deliver the stimulations to the insect, a short length of column at the output of the transfer line was inserted into the small hole in the glass chamber used in the EAG3 experiments.

We manually injected 1µL of solution, which consisted of hexane loaded with a blend of synthetic compounds diluted at 10 ng.µL^-1^ or 1 µg.µL^-1^ each, into the GC. In total, 37 synthetic compounds (Table S3) were split into two blends, which were delivered to the same individual in a random order. One run lasted 30 minutes, i.e. one experiment lasted 1 hour. Negative (mineral oil) and positive (hexenyl acetate and 1-octen-3-ol, dilution v/v: 10^-2^ in mineral oil) control stimulations were successively performed before and after each run, following the same procedures as for the EAG3 experiments.

### Neuronal model of olfactory detection

Our modelling approach was designed to formally validate the two following hypotheses. (1) The detection hypothesis: for a given number of olfactory receptors (ORs), the polyphagous species’ ecological need to detect hosts selects stronger olfactory responses to shared volatiles compared to species-specific host volatiles. (2) The alternative discrimination hypothesis: the polyphagous species’ use of volatiles to discriminate between hosts selects stronger olfactory response to species-specific compounds compared to shared host compounds. The modelling workflow was as follows: we designed a large number of simple neuronal models of insect olfactory systems, each one responded to a random selection of compounds with random sensitivity. This set of models is intended to include all possible combinations that may have been reached through random mutations, although only some are realistic under natural conditions. We then identified the subset of models under hypothetical selection and analysed their specific properties. To do this, we fed samples of intact or sliced fruit volatile emissions into the model (see collection method above). By drawing on these data sets, we estimated how effectively each model detected or discriminated between fruit species.

More specifically, each model included three categories of virtual compartments: compounds (n = 665 or 511, from sliced and intact fruit, respectively), ORs (one OR unit in the model includes virtually all the olfactory sensory neurons that express the same OR), and total antennal activity (equivalent to EAG3). An odour (input) was defined by a combination of compounds at different doses (in log units). Each OR was activated by a fixed number of compounds whose identity was randomly chosen. Thus, while some compounds did not activate any OR, others activated several ORs. A compound is considered to have been detected if it activates at least one OR. Each compound dose/OR response relationship was defined by two parameters: a sensitivity threshold (drawn from a uniform distribution covering 5 log units) and a dynamic range width (drawn from a log-normal distribution of mean 3 and standard deviation 1). The activation xij of an ORi by compound j was 0 below threshold, 1 above threshold + dynamic range width, and increased linearly between the two. The activity of ORi was calculated by combining all compound activations with the following equation: 1 – ᴨ_j_(1-x_ij_), which ensured that the activity level remains between 0 and 1. The antennal activity (EAG) was the sum of OR activities.

For each model, a fruit detectability index, which estimates the effectiveness of the insect sensory system to detect fruit, was defined as the sum of total antennal response to all fruit odours. In order to estimate the ease with which the sensory system discriminates fruit species, we calculated a fruit species discriminability index as follows. A random forest analysis was applied to classify the patterns of OR responses to volatile samples into the different fruit species (randomForest function, R package randomForest). The index was defined as the opposite of the out-of-bag scores, i.e. the number of correctly classified samples divided by the total number of samples (Figure 3B). The random forest classifier is thus considered as a proxy of the brain’s ability to discriminate fruit on the basis of OR responses. We also calculated a joint functionality index, defined as the product of the fruit detectability and the fruit species discriminability indices, each centred on 0.5 and with standard deviation adjusted for values between 0 and 1.

We then built 12 sets of 10,000 models. Each model had a combination of a fixed number of ORs (10 or 20, a rough estimate of the number of ORs likely to be sensitive to fruit odours in Tephritidae species), and a fixed number of compounds detected per OR (low, medium or high value, i.e. 10, 20 or 60 compounds per OR for models with 10 ORs, 5, 10 or 30 compounds per OR for models with 20 ORs, respectively). Volatile data for sliced or intact fruit was fed into the models.

### Behavioural assay setup

An *ex situ* trapping system was used to study the behavioural response of *B. dorsalis* to different blends of compounds. The system consisted of two handmade traps, which were placed in a rearing cage measuring 30 cm x 30 cm x 30 cm. The traps were made using a 30 mL wide mouth glass vial sealed with parafilm and lined with yellow coloured tape to guide the flies towards the traps. The lid was then perforated and a sectioned transparent 1,000 µL pipette tip was inserted into the hole, serving as a conduit for the flies. A piece of filter paper loaded with 1 µL of the tested blend was placed at the bottom of the trap and the odour diffused passively. The two traps in the cage were loaded either with different blends or with a negative control (mineral oil only). Six cages were tested simultaneously with the same combination of traps in different positions. In each cage, 30 starved gravid female flies were released simultaneously and the number of flies in each trap was noted every 30 or 60 minutes. Each trap was used only once and the cages were thoroughly cleaned with soap between experiments.

For behavioural assays, two blends of 5 compounds were tested. The first blend consisted of methyl 2-methylbutyrate, butyl butyrate, methyl heptanoate, methyl octanoate and prenyl acetate. The second blend consisted of ethyl acetate, cis-3-hexenyl acetate, hexyl acetate, ethyl propionate and ethyl butyrate. Lastly, we used a blend combining all ten compounds. Preliminary work for calibrating compound ratios was carried out in the SLU laboratory at the Swedish University of Agricultural Sciences in order to homogenize the volatile emission rates in the blend before the behavioural assays. The blends were diluted in mineral oil and tested at two different doses (dilution v/v 10^-4^ and 10^-3^ of total compounds). The percentage of captured flies was analysed using generalized linear mixed-effects models (GLMM) with binomial error distribution (*glmer* function, R package *lme4*). Blends and doses were considered as fixed effects and the experimental date was considered a random effect.

### Data analysis

A pre-processing of GC-MS data was carried out with MZmine2 software (Pluskal et al., 2010). Compound peaks were detected by ADAP algorithm (Ni et al., 2016) (S/N threshold 3, Wavelet Coeff. 3, minimum feature height 1,000,000, coefficient/area threshold 300, peak duration 0 to 0.2, RT wavelet 0 to 0.05 min), deconvoluted through blind source separation (window width 0.1, minimum number peaks 2, RT tolerance 0.025), RANSAC aligned (m/z tolerance 0.3, RT tolerance 0.1 min, automatic iteration no., minimum points no. 50%, threshold 0.1 min), and gap filled (m/z tolerance 0.3, RT tolerance 0.02 min). Each compound was identified according to its molecular weight and retention index (RI, calculated from the retention times of C6-C30 n-alkanes), and by comparing its mass spectra with NIST 14 library databases.

Further statistical analyses were conducted using R software (R Core Team, 2020). When the quantity of compounds in the control samples was greater than in the samples with fruit, they were excluded from the analysis (p < 0.05, Wilcoxon test, *wilcox.test* function, R package *Stats*). Volatilomics data were then normalized by dividing each compound peak by the geometric mean of each corresponding sample, then Box-Cox transformed (*boxcox* function, R package *MASS*, lambda = 0.014) and Pareto scaled (*paretoscale* function, R package *RFmarkerDetector* (van den Berg et al., 2006; Grace and Hudson, 2016)).

The degree of sharedness among fruit species was calculated for each compound by Shannon α-diversity index (*diversity* function, R package *vegan*) (Jost, 2007). For the calculation, we used the volatile samples of intact (α-div_IF_28) and sliced (α-div_SF_ ^28^) fruit of 28 species, as well as volatile samples of intact (α-div_IF_^13^) and sliced (α-div_SF_ ^13^) fruit from a subset of 13 species. The subset included species with a *B. dorsalis* infestation rate of above 5%, which were collected in Réunion during a recent campaign to monitor Tephritidae infestation (Table 1) (Moquet et al., 2021). The indices α-div_IF_28 and α-div_SF_ ^28^ can vary between a minimal value of 0 (if a single fruit species emits the compound) and a maximal value of 3.33 (when all 28 fruit species emit the compound). Similarly, the indices α-div_IF_^13^ and α-div_SF_^13^ can vary between a minimal value of 0 (when a single fruit species emits the compound) and a maximal value of 2.57 (when all 13 fruit species emit the compound). To measure the phylogenetic signal, Abouheif’s *C_mean_* index (*abouheif.moran* function, R package *adephylo*) was computed for each compound based on an autocorrelation approach (Abouheif, 1999; Pavoine et al., 2008). To calculate the index, the phylogeny of the 28 fruit species was generated by Bayesian phylogenetic inference using Markov Chain Monte Carlo (MCMC) methods (Ronquist et al., 2012). The matrix of phylogenetic proximities was performed with the “oriAbouheif” method (*proxTips* function, R package *adephylo*) (Pavoine et al., 2008). It was calculated for the 28 species using volatile samples of intact (C ^28^) and sliced (C ^28^) fruit.

After a preliminary demodulation of the EAD3 signal at the chopping frequency (1 Hz), we calculated the total antennal activity from EAG3 and EAD3 data using the current source density (CSD) model (Jacob, 2018; Ramiaranjatovo et al., 2023). To apply the linear models, antennal activity data were normalized by adding a constant value before applying a Box-Cox power transformation. The constant value was chosen to ensure that the Box-Cox transformation was log-equivalent. The normality of the resulting distribution was assessed using a Shapiro-Wilk test. For linear models, individual identity was used as an independent variable.

## Data availability

Data have been deposited in a long term repository service : Jacob, Vincent, 2024, “Raw data and code for Ramiaranjatovo et al. (2024) “Olfaction in Tephritidae: a balance between detection and discrimination””, https://doi.org/10.18167/DVN1/T3XUXW, CIRAD Dataverse, DRAFT VERSION and will be available upon publication.

## Code availability

The code used for modelling has been deposited in a long term repository service : Jacob, Vincent, 2024, “Raw data and code for Ramiaranjatovo et al. (2024) “Olfaction in Tephritidae: a balance between detection and discrimination””, https://doi.org/10.18167/DVN1/T3XUXW, CIRAD Dataverse, DRAFT VERSION and will be available upon publication.

## Acknowledgements

The authors would like to thank Serge Glenac and Jim Payet for collecting and rearing the insects. The authors greatly acknowledge the Plant Protection Platform for their technical support (3 P, IBISA). This research was conducted within the frameworks of the research platform ‘Biocontrôle et épidémio-surveillance végétale en ocean indien’ (https://www.dp-biocontrole-oi. org/) and the UMT BAT ‘Biocontrôle en Agriculture Tropicale’. This work was funded by the Conseil Régional de la Réunion and the European Regional Development Fund (ERDF), the Centre de Coopération Internationale en Recherche Agronomique pour le Développement (CIRAD), and the French Ministry of Agriculture, as part of the research project GEMDOTIS (Ecophyto II 2018) and ATTRACTIS (Ecophyto II 2023).

---

**Figure S1.**
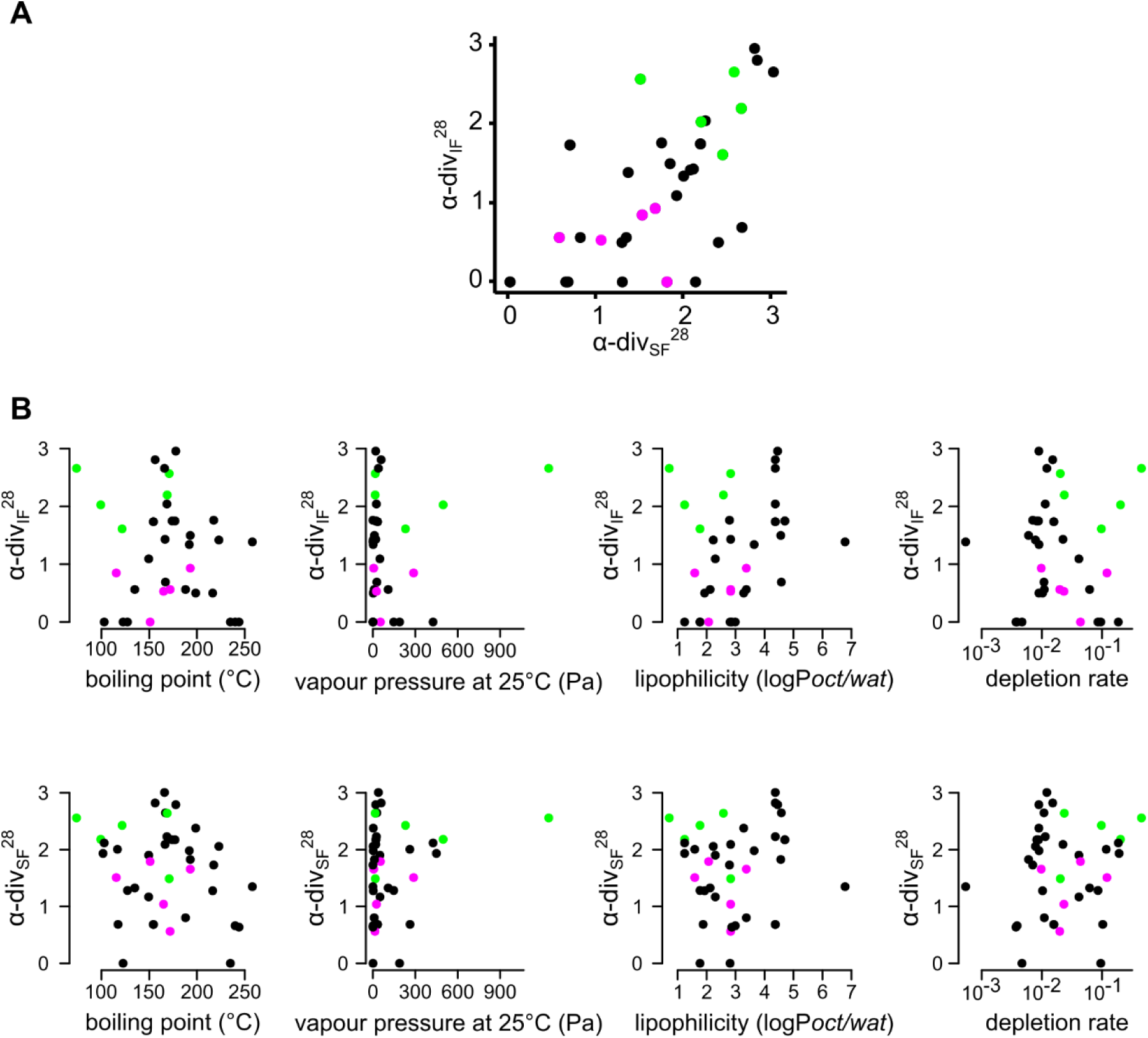
Chemical characteristics of the 40 compounds used for testing the olfactory system. 30 of these compounds were used for EAG3 tests and 37 were used for GC-EAD3 tests. (A) Relationship between the fruit α-diversity indices calculated from intact and sliced fruit. Coloured dots show the compounds used for behavioural tests: the five compounds included in the species-specific compound blend are in magenta and the five compounds included in the shared fruit compound blend are in green. (B) Indices of α-diversity are not correlated with any chemical properties. Same conventions as in panel (A).

**Figure S2.**
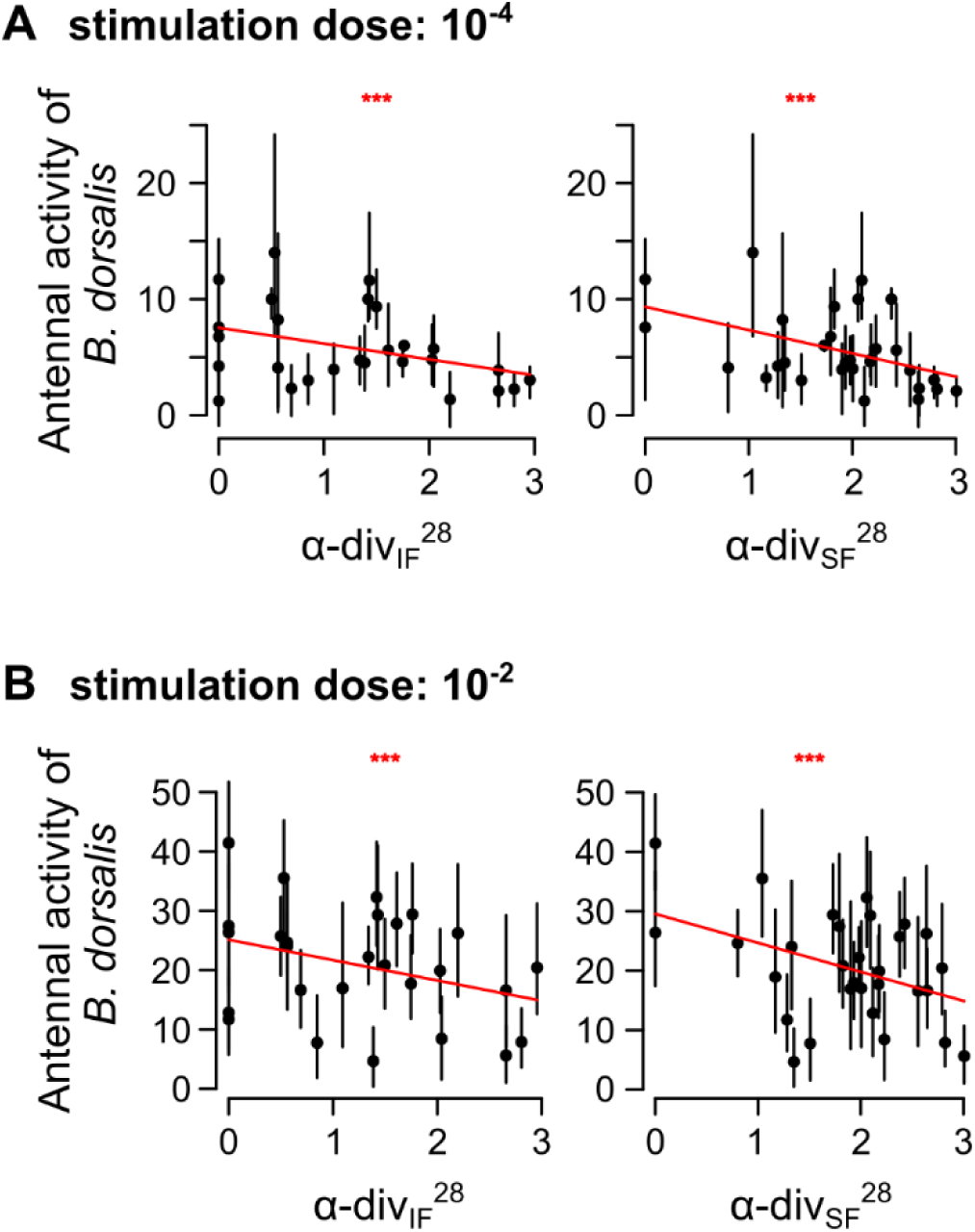
Antennal responses of *Bactrocera dorsalis* measured with EAG3 correlate with fruit α-diversity calculated from intact and sliced fruit emissions. (A-B) Scatterplot at the two doses tested. Same conventions as in Figure 4.

**Figure S3.**
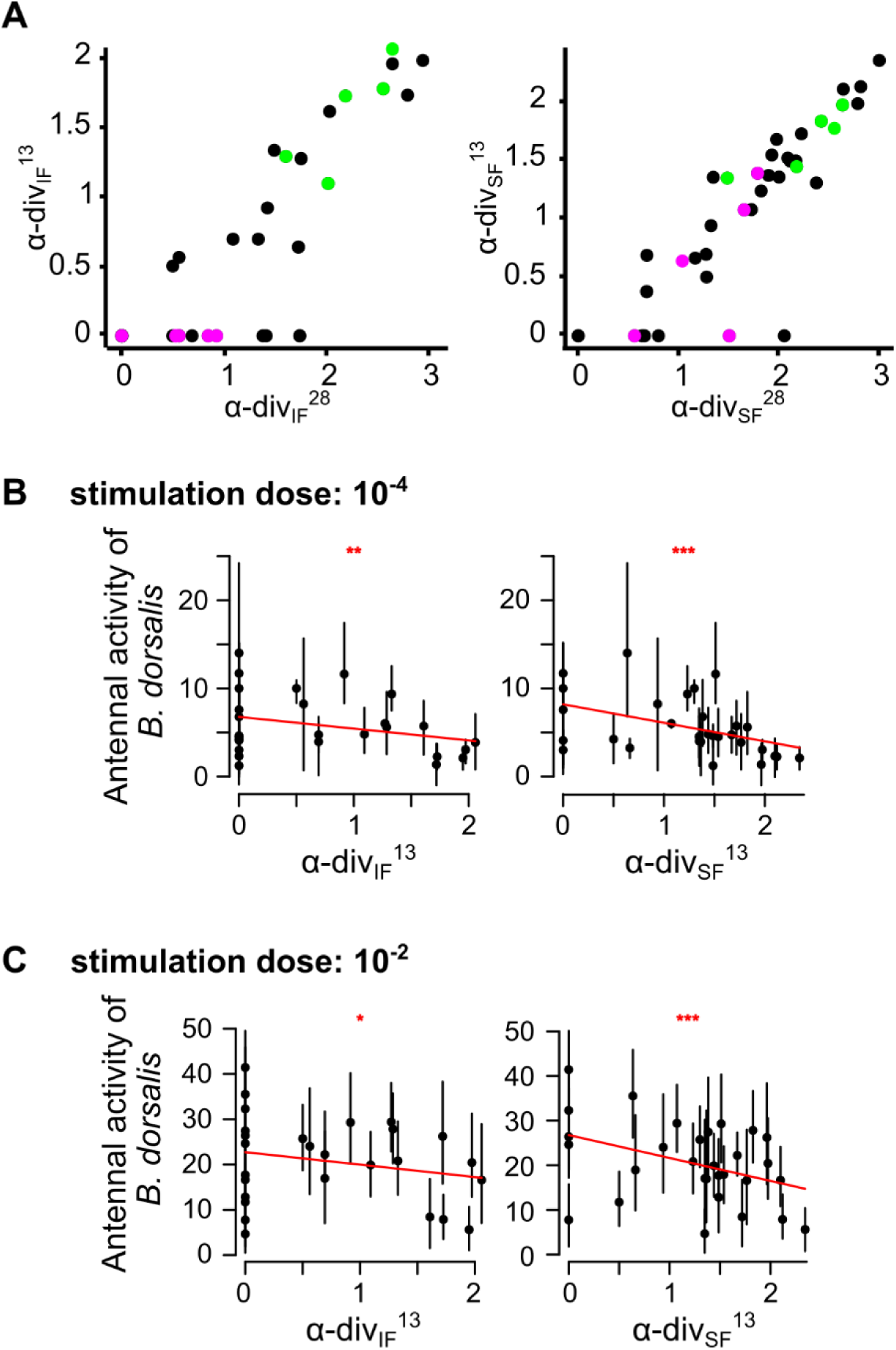
Antennal responses of *Bactrocera dorsalis* measured with EAG3 correlate with indices of α-diversity. Indices of α-diversity were calculated from a subset of 13 intact or sliced fruit emissions (Table 1), at the two doses tested. (A) relationship between α-diversity indices calculated from 13 of the 28 fruit species. (B-C) Scatterplots at the two dose tested. Same conventions as in Figure S1 and S2.

**Figure S4.**
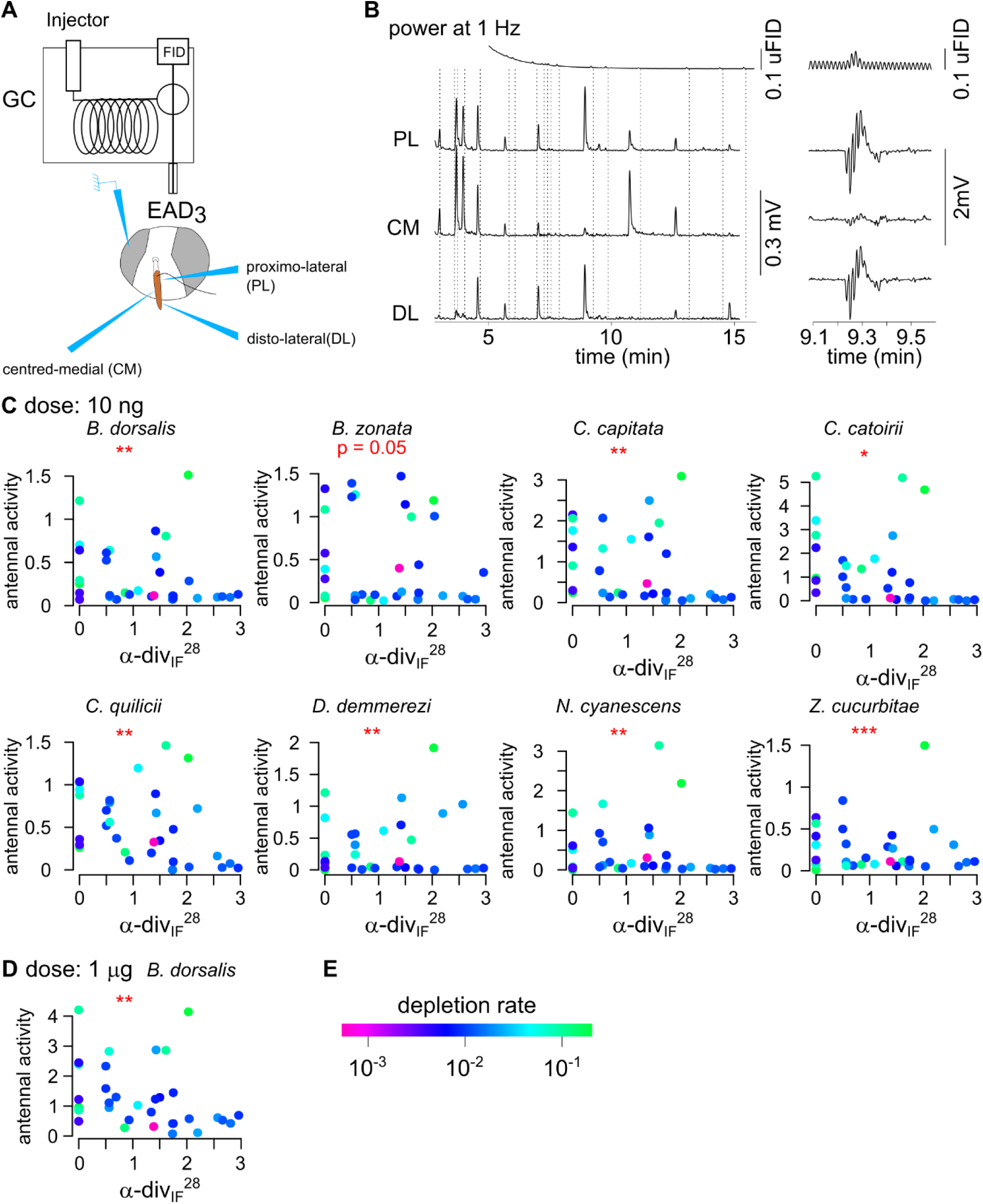
Antennal responses of eight Tephritidae species in GC-EAD3. (A) Schematic representation of the chopped GC-EAD3 system. (B) An example of GC-EAD3 recording on individual female *B. dorsalis*. Left: Signal is demodulated at 1 Hz, the frequency of the chopping modulation. The first line is the FID signal and the three lines below are the EAD signals at the three antennal positions. Right: zoom on a single response, showing the full signal before demodulation. (C-D) Antennal responses to synthetic compounds at dose 10 ng (C) and 1 µg (D, n=4), depending on the fruit α-diversity of intact fruit emissions. Each point represents a mean response to a compound (n = 4 for dose 10 ng, n = 5 for dose 1 µg). (E) Compound depletion rate is colour coded. The significance of a correlation between antennal activity and an interaction between α-div_IF_28 and depletion rate is indicated (* p < 0.05, ** p < 0.01, *** p < 0.001).

**Figure S5.**
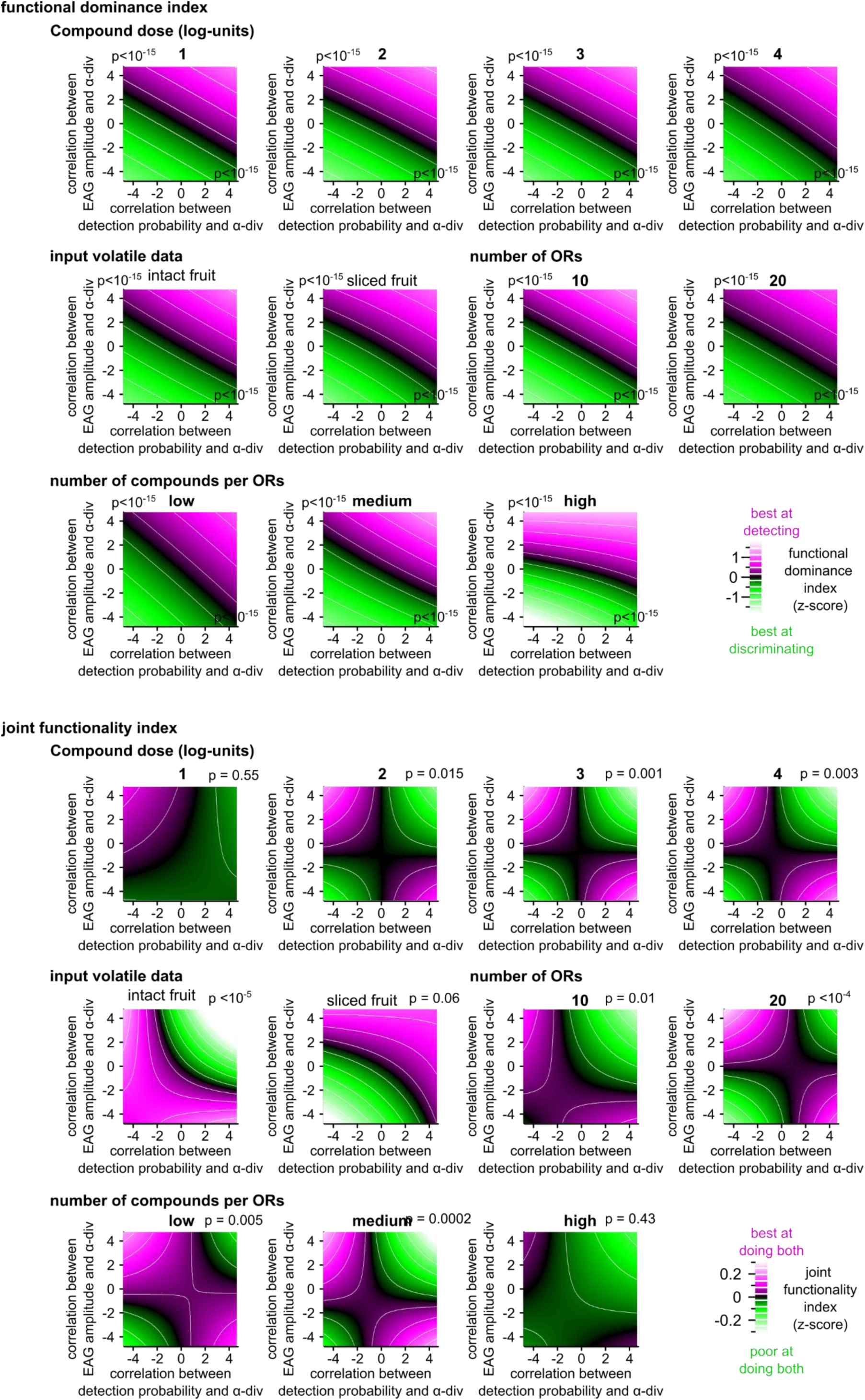
Functional dominance index (top graphs) and fruit detectability index (bottom graphs) dependence on EAG response properties, as in Figure 4, depicted for all stimulation dose, input volatile data, number of ORs in the model, and number of compounds per ORs in the model. The functional dominance index is simply the difference between detectability and discriminability indices. The p-values indicate how much it depends on the two axis in each case (linear model). For joint functionality index, the p-values indicate how much it depends on an interaction between the two axis.

**Figure S6.**
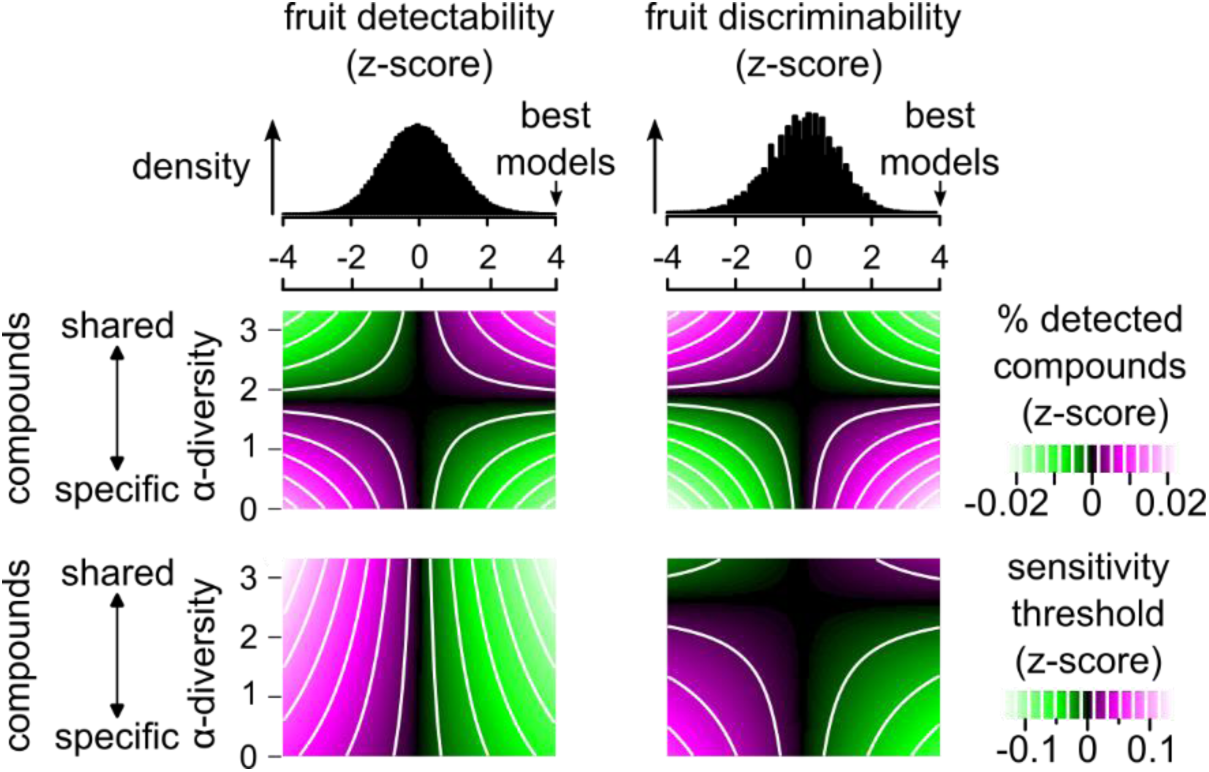
Model parameters for 120,000 models with random connectivity. Top: the density distribution of fruit detectability and the fruit-species discriminability index for the 120,000 models. The best models for the two functions are on the upper part of the distribution. The linear projection of the proportion of compounds detected by the olfactory system (upper heatmaps) and that of the sensitivity threshold of detected compounds (lower heatmaps) are drawn on the spaces delineated by the model’s fruit detectability indices (left heatmaps) and fruit-species discriminability indices (right heatmaps), as the x-axis, and compound α-diversity as the y-axis. that response parameters. The proportion of detected compounds, the sensitivity thresholds of detected compounds and, to a lesser extent, the dynamic range of detected compounds depended on a joint effect between compound α-diversity and the fruit detectability index (F (1, df > 10^7^) = 285; p < 10^-15^ ; F (1, df > 10^7^) = 613; p < 10^-15^ ; and F (1, df > 10^7^) = 34.3; p < 10^-8^, respectively). Specifically, olfactory systems that are efficient at detecting fruit perceive shared compounds better than species-specific ones. Inversely, the same response parameters depended on a joint effect between fruit α-diversity and fruit species discriminability index, namely the proportion of detected compounds (F (1, df > 107) = 453; p < 10-15), the sensitivity thresholds of detected compounds (F (1, df > 107) = 731; p < 10-15) and, to a lesser extent, the dynamic range of detected compounds (F (1, df > 107) = 18.6; p < 10-4). Olfactory systems capable of discriminating between fruit species efficiently are better at detecting species-specific compounds than shared compounds.

**Table S1.** List of names, classes, α-div^28^, α-div^13^ and C_mean_ values for the 511 compounds from intact fruit samples

**Table S2.** List of names, classes, α-div^28^, α-div^13^ and C_mean_ values for the 665 compounds from sliced fruit samples.

**Table S3.**
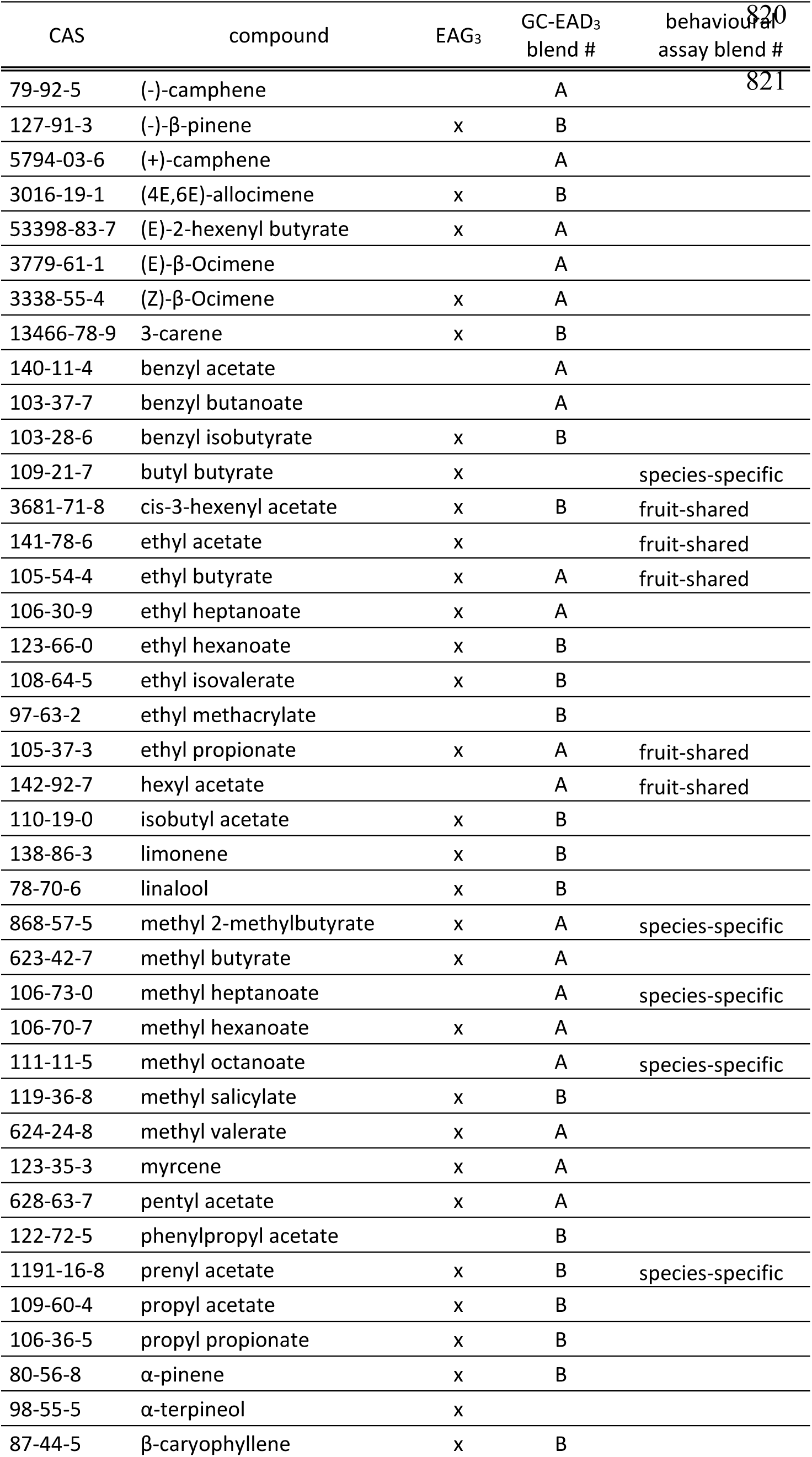
List of synthetic compounds used for electrophysiological and behavioural tests. Compounds were asigned to two different blends for EAD and behavioural tests.

